# DNA Methylation Associated with Postpartum Depressive Symptoms Overlaps Findings from a Genome-wide Association Meta-Analysis of Depression

**DOI:** 10.1101/600742

**Authors:** Dana M. Lapato, Roxann Roberson-Nay, Robert M. Kirkpatrick, Bradley T. Webb, Timothy P. York, Patricia Kinser

## Abstract

**Background:** Perinatal depressive symptoms have been linked to adverse maternal and infant health outcomes. The etiology associated with perinatal depressive psychopathology is poorly understood, but accumulating evidence suggests that understanding inter-individual differences in DNA methylation (DNAm) patterning may provide insight regarding the genomic regions salient to the risk liability of perinatal depressive psychopathology.

**Results:** Genome-wide DNAm was measured in maternal peripheral blood using the Infinium MethylationEPIC microarray. Ninety-two participants (46% African-American) had DNAm samples that passed all quality control metrics, and all participants were within seven months of delivery. Linear models were constructed to identify differentially methylated sites and regions, and permutation testing was utilized to assess significance. Differentially methylated regions (DMRs) were defined as genomic regions of consistent DNAm change with at least two probes within 1kb of each other. Maternal age, current smoking status, estimated cell-type proportions, ancestry-relevant principal components, days since delivery, and chip position served as covariates to adjust for technical and biological factors. Current postpartum depressive symptoms were measured using the Edinburgh Postnatal Depression Scale. Ninety-eight DMRs were significant (False Discovery Rate < 5%) and overlapped 92 genes. Synaptic signaling, neural development, and platelet formation were the most represented biological processes in gene set analysis, and comparison to the 44 loci discovered in the latest Psychiatric Genomics Consortium meta-analysis of depression revealed 3 overlapping regions and significant enrichment (p<0.03).

**Conclusions:** Many of the genes identified in this analysis corroborate previous allelic, transcriptomic, and DNAm association results related to depressive phenotypes. Future work should integrate data from multi-omic platforms to understand the functional relevance of these DMRs and refine DNAm association results by limiting phenotypic heterogeneity and clarifying if DNAm differences relate to the timing of onset, severity, duration of perinatal mental health outcomes of the current pregnancy or to previous history of depressive psychopathology.

## Introduction

Perinatal depressive symptoms can occur any time during pregnancy or shortly following birth and have been associated with an increased risk for episodes of major depression with onset in the peripartum (MDP), pregnancy complications, maternal suicide, and adverse infant health outcomes and development.^1–7^ The Diagnostic and Statistical Manual (5^th^ edition; DSM-5) classifies MDP as a major depressive episode that occurs during pregnancy or within 4 weeks of delivery;^8^ however, in practice, researchers and clinicians may extend this period to up to one year postpartum. Attempts to understand the impact of depressive psychopathology on biological mechanisms have primarily centered on elucidating the relationship between maternal mental health and negative infant outcomes. As a result, relatively little is known regarding how perinatal depressive symptoms affect maternal biological processes. Epidemiological studies suggest that episodes of major depression (MD) may increase the risk for other adverse health outcomes, such as cardiovascular disease, and perturb immune system activities. The biological pathways associated with these persistent changes in immune activity and elevated risk have not yet been identified. More work is needed to uncover the pathways underlying these comorbidities and to determine if the biology impact from depressive symptoms differs by clinical subtype (e.g., perinatal, early-onset, etc.). One potential avenue for understanding biological changes associated with depressive psychopathology is through DNA methylation (DNAm) studies.

DNAm is a chemical modification typically found on cytosines bordering guanines (i.e., cytosine-phosphate-guanine [CpG] sites). DNAm can influence gene expression, genomic stability, and chromatin conformation.^9^ Inter-individual differences in DNAm have been associated with early mortality,^10^ cancer,^11^ imprinting disorders,^12,13^ childhood trauma exposure,^14^ biological age,^15,16^ and schizophrenia.^17,18^ Associations between DNAm and clinical MD and/or depressive symptoms proximal to^19–22^ or absent pregnancy^23–26^ have been reported; however, most of the perinatal depression studies have focused on identifying DNAm patterns in fetal tissues associated with maternal mental health, leaving a significant knowledge gap. Characterizing the relationship between DNAm and perinatal depressive symptoms may provide insight regarding the pathoetiology of MDP as well as potentially identify biological markers.^27^

This study sought to identify DNAm patterns associated with perinatal depressive symptoms during the first seven months postpartum in maternal blood focusing on regional DNAm changes. Genome-wide DNAm and repeated measures of maternal mental health were collected as part of a longitudinal study of preterm birth.^28^ The rationale for focusing on differentially methylated regions (DMRs) rather than single CpG site associations was twofold. One, regional changes are thought to represent differences more likely to be biologically meaningful and statistically credible.^29^ A single CpG site associated with a trait could be spurious; however, multiple CpG sites within one region each associating with a trait in the same direction is likely to represent a more robust finding. Two, regional analyses reduce the burden of multiple tests and allow one to test for smaller probe effect sizes.^30^ The sample size for this study is acceptable to test regional differences, but it is not well-powered to identify individual probe associations.

## Methods

### Study participants

The data for this analysis come from the Pregnancy, Race, Environment, Genes (PREG) study and its postpartum extension.^28^ Both PREG and the extension received IRB approval, and all participants provided written informed consent for both parts. Most participants had the first postpartum visit within three months of delivery and the second visit within nine months.

### Study eligibility criteria

Participants were required to meet the same enrollment and birth inclusion criteria used in the PREG study.^28^ In summary, enrollment criteria required participants to 1) be <21 weeks gestation, 2) have a singleton pregnancy, 3) have not used artificial reproductive technology for the current pregnancy, 4) be between 18 and 40 years old, and 5) be absent of major health conditions (e.g., diabetes). Additionally, the participant and the biological father had to self-identify as either both African-American or both European-American and without Middle Eastern or Hispanic ancestry. Birth exclusion criteria included chromosomal abnormality, polyhydramnios/ oligohydramnios, placenta previa, and congenital birth defects.

### Psychiatric assessments

Current perinatal depressive symptoms were measured at both postpartum study visits using the Edinburgh Postnatal Depression Scale (EPDS).^31^ The EPDS is frequently used to assess perinatal depression because the items focus on symptoms specifically related to depression that would not be part of a typical pregnancy.^32^ For example, difficulty sleeping is common in pregnancies regardless of maternal MD status. EPDS total score was kept continuous.

### Genome-wide DNAm measurement and processing

Maternal DNAm was assayed from peripheral blood using the Illumina EPIC beadchip, which includes more than 850,000 probes and interrogates regulatory, genic, and intergenic regions.^33^ Blood samples were collected at each study visit along with health questionnaires. DNA was extracted at Virginia Commonwealth University and sent to HudsonAlpha Laboratories for DNAm measurement. Peripheral blood was selected given its accessibility, the availability of cell-type deconvolution methods, and the strong evidence of immune system involvement in MDP pathophysiology.^34–37^

Raw microarray data was processed using Bioconductor packages in the R environment in line with best practices^38–40^ (version 3.5). Briefly, signal intensity and probe failure rate were evaluated using the *minfi*^41^ package to identify poor quality samples and probes. Samples were removed if either the median unmethylated or methylated signal intensities were less than 10.5. Probes were removed if they failed in >1% of samples (n=12,557), overlapped single nucleotide polymorphisms (n=30,435), or had been identified as cross-hybridizing in the Illumina Infinium HumanMethylation450 beadchip (predecessor technology).^42^ Probes on the sex chromosomes were retained given that the entire sample was female, leaving a total of 782,884 probes. All samples were quantile normalized, and blood cell type proportions were estimated using the Houseman method.^43^ Sample identity was confirmed using the 59 control probes on the EPIC beadchip. These probes overlap polymorphic sites and in aggregate can estimate sample relatedness and detect sample duplication. All pairwise sample correlations were calculated. Any sample correlated too poorly with its sister samples (r < .8) or too highly with samples from another person (r > .6) were removed or relabeled if the correct identity could be ascertained. Only one blood sample per person was used for this analysis. In general, DNAm samples from the first postpartum visit were used; however, if a participant’s first postpartum visit failed quality control and she had a second postpartum DNAm sample within seven months of delivery, then that sample and the EPDS questionnaire from the same visit was used. The purpose for using only one sample came from a desire to limit phenotypic heterogeneity. Perinatal depressive psychopathology is not fully understood, but meta-analysis suggests that there may be something phenotypically unique about depression during the first few months postpartum.^44^ Point estimates of major and minor episodes of MDP are highest in the third month postpartum (∽13%) and drop noticeably after the seventh month postpartum (∽6.5%).^45,46^ Limiting the study to include only samples from the seven months postpartum captured the time period with the highest estimates of postpartum depressive symptom prevalence and reduced the phenotypic heterogeneity.

## DNAm Analysis

### Covariate selection

Principal component analysis (PCA) was applied to the normalized methyl values^47^ of the filtered probe set, and correlations between the top ten principal components (PC) and technical and biological variables were plotted to identify potential confounders. Using this information, microarray row, granulocyte proportion, and median unmethylated signal intensity were selected as covariates in addition to maternal age, days postpartum at blood collection, and smoking status, which were selected *a priori* based on known or putative associations with DNAm. Slide effects were addressed using ComBat.^48^ Allelic ancestry was assessed using the methods described in Barfield et al. (2014).^49^ Briefly, PCA was applied to probes that directly overlapped single nucleotide polymorphisms (i.e., CpG-SNPs), and the top ten PCAs from that analysis were correlated with self-reported race to identify the PCs that most strongly associated with race. Two allelic PCs were selected as covariates based on their relationship with self-reported race (see Supplementary Figure at https://osf.io/v7aez).

### Identifying single site and regional DNAm changes

The methyl values for individual probes were regressed onto postpartum EPDS total score using the *limma* package.^50^ Covariates controlling for row, granulocyte proportion, median unmethylated signal intensity maternal age, gestational age at blood collection, smoking status, and allelic ancestry were included. The strength of the association between individual probes and EPDS total score was evaluated using empirical p values derived from k=20,000 permutations. The median effect size for probes used in the DMR analysis was assessed using the difference in adjusted R^2^ values between the full model and a reduced model without EPDS total score.

For regional analysis, the probes with the largest observed t statistics (top and bottom 2.5% of tested probes) were considered. Differentially methylated regions (DMRs) were defined as contiguous regions of consistent DNAm change (i.e., all hypermethylated or all hypomethylated) that contained at least two probes within 1kb using a method similar to that described by Ong and Holbrook.^30^ This strategy is similar to the DMRcate algorithm in that only the subset of the probes with the best evidence for association is used to construct DMRs.^51^

DMR significance was assessed using a permutation strategy. DMRs were constructed from both the observed data and 1000 of the 20,000 DMP permutations. For each DMR, the area under the curve (AUC) was calculated using the trapezoidal rule, where each probe’s t statistic served as height and the distance between the probes as width. In this way, the magnitude of the AUC represents both the strength of each probe’s association (height) and the size of the region (width). The Significance of Analysis Microarray (SAM) method was implemented to assign test statistics to each observed DMR.^52^ This method ranks all DMRs generated within a permutation by AUC. Row-wise comparisons between the ranked observed DMRs and the ranked permutation DMRs is used to calculate the false discovery rate (FDR).

### Gene set enrichment and comparison to other genetic findings related to depression

DMPs and DMRs were annotated using AnnotationHub.^53^ Gene set enrichment testing for functional and regulatory roles was performed on the combined dataset of DMPs and DMRs using Entrez IDs. The rationale for combining DMRs and DMPs into a single group for gene set analysis was to address the issue that not all probes are capable of forming DMRs. In order to give those regions of the genome an opportunity to contribute to gene set enrichment analysis, DMPs and DMRs were analyzed together (see OSF Supplement for DMP-only and DMR-only enrichment analyses).

The results from this analysis were compared directly to two studies of depression: the latest Psychiatric Genomics Consortium (PGC) meta-analysis of genome-wide association studies of depression and an epigenome-wide association study (EWAS) of early-onset MD. For the PGC study, the 44 significant loci were obtained to determine the extent of overlap with significant DNAm regions.^54^ Bootstrap and permutation methods (k = 1000) were used to test if DNAm regions were enriched for PGC loci. For the early-onset MD EWAS, site, regional, and gene enrichment results from the Adolescent and Young Adult Twin Study (AYATS) were compared to findings from this study to determine the extent of overlap and similarity.^26,55^

## Results

### Sample characteristics

Sample demographics can be found in Table 1 and are representative of Richmond, Virginia. Approximately half of the participants (46%) self-identified as African-American. Most of the women (65%) were primigravida and had full-term pregnancies (94%). Very few were current smokers (8%), and 18% of the total sample self-reported a positive lifetime history of MD. Both the blood sample for DNAm measurement and the health questionnaire (including the EPDS) were collected during this study visit. The average time between delivery and postpartum study visit was 57 days.

**Table 1:**
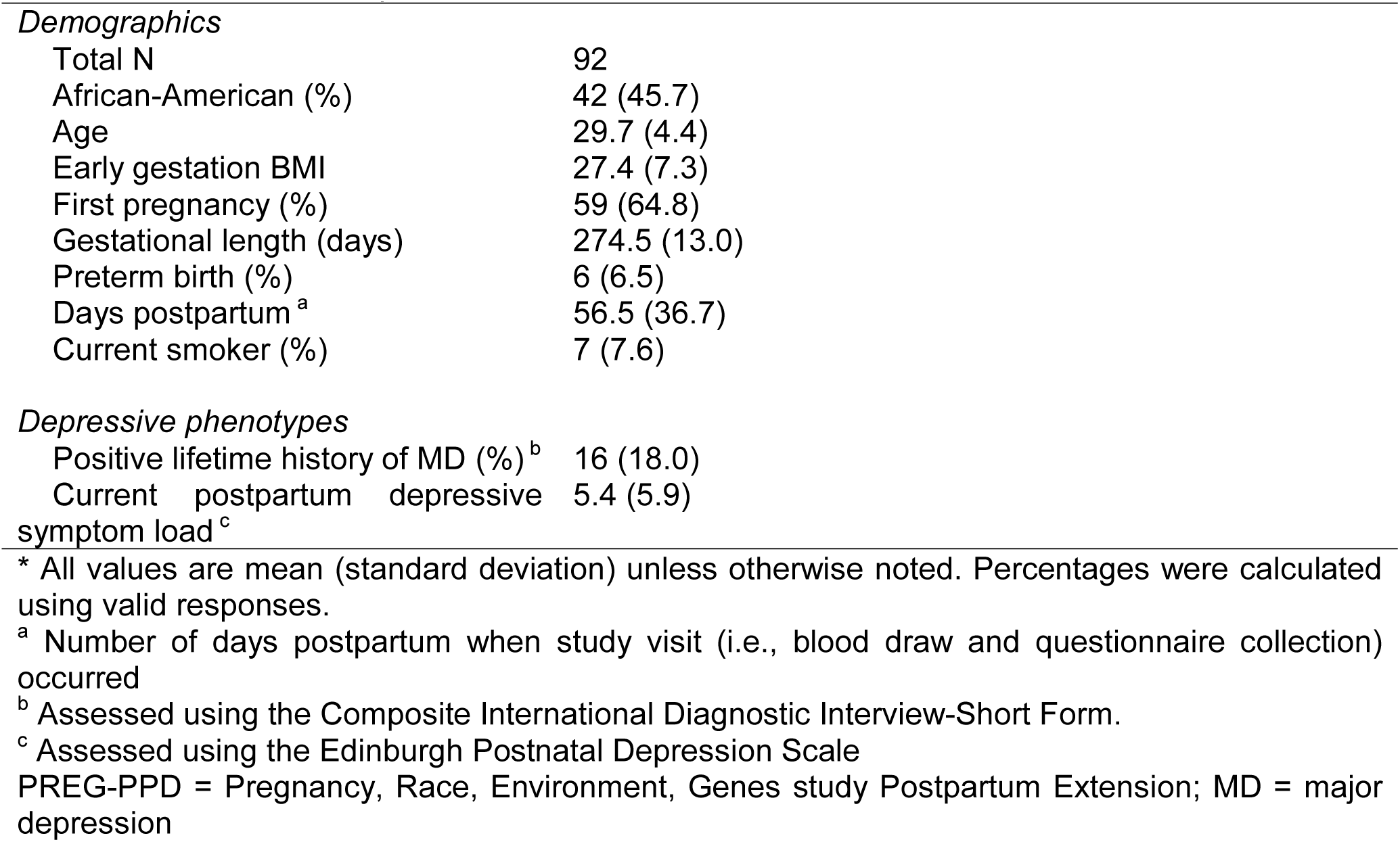
PREG-PPD Sample Characteristics

|  |  |
| --- | --- |
| Total N | 92 |
| African-American (%) | 42 (45.7) |
| Age | 29.7 (4.4) |
| Early gestation BMI | 27.4 (7.3) |
| First pregnancy (%) | 59 (64.8) |
| Gestational length (days) | 274.5 (13.0) |
| Preterm birth (%) | 6 (6.5) |
| Days postpartum <sup>a</sup> | 56.5 (36.7) |
| Current smoker (%) | 7 (7.6) |
| *Depressive phenotypes* |  |
| Positive lifetime history of MD (%) <sup>b</sup> | 16 (18.0) |
| Current postpartum depressive symptom load <sup>c</sup> | 5.4 (5.9) |

\* All values are mean (standard deviation) unless otherwise noted. Percentages were calculated using valid responses.
<sup>a</sup> Number of days postpartum when study visit (i.e., blood draw and questionnaire collection) occurred
<sup>b</sup> Assessed using the Composite International Diagnostic Interview-Short Form.
<sup>c</sup> Assessed using the Edinburgh Postnatal Depression Scale
PREG-PPD = Pregnancy, Race, Environment, Genes study Postpartum Extension; MD = major depression

### Differentially methylated probes and regions

After microarray quality control and processing, 782,884 probes remained for analysis. From this filtered probe set, it was possible to create up to 109,340 background DMRs. N=206,804 probes were ineligible to participate in regional analyses because they did not have a neighboring probe within 1kb. Individual site analysis of the entire probe set identified 50 DMPs significantly associated with EPDS total score (empirical p = 0 after 20,000 permutations; see https://osf.io/q2s8d for summary statistics). Approximately 39,150 probes were taken forward for regional analysis. The median difference in adjusted R^2^ values for the full versus reduced models for these probes was .053 (interquartile range = .02) Ninety-eight genomic regions spanning a collective 116.2kb were significantly differentially methylated by postpartum depressive symptom load (FDR < 5%). The significant regions overlapped 92 genes on 20 chromosomes (none on chromosomes 7 or 18), and on average spanned ∽1.2kb in length (see https://osf.io/bf6sw for complete gene list). The number of CpG probes in significant DMRs ranged from 2 to 10 (mean = 3.48 probes).

### Gene set enrichment and comparison of DNAm patterns associated with postpartum depressive symptoms and other genetic findings related to depression

Detailed results for the combined gene set enrichment of DMPs and DMRs can be found in Table 2. In short, the combined analysis identified one biological process (BP; cognition) and four cellular components (CC; DNA repair complex, neuron to neuron synapse, axon part, and synapse part) significant at an FDR < 5%. The DMP and DMR only analyses did not identify any categories with an FDR < 5% and produced dissimilar results (see supplement for DMP-only and DMR-only tabular results). The DMP only analysis identified multiple BP and CC categories associated with neural processes (e.g., central nervous system neuron development, transmission of nerve impulses, somatodendritic compartment, neuronal cell body). The DMR only analysis returned Gene Ontology (GO) categories from a variety of biological systems, including platelet formation and morphogenesis, cardiac muscle tissue development, chemical synaptic transmission, inflammatory cell apoptotic process, and tissue morphogenesis.

**Table 2:**
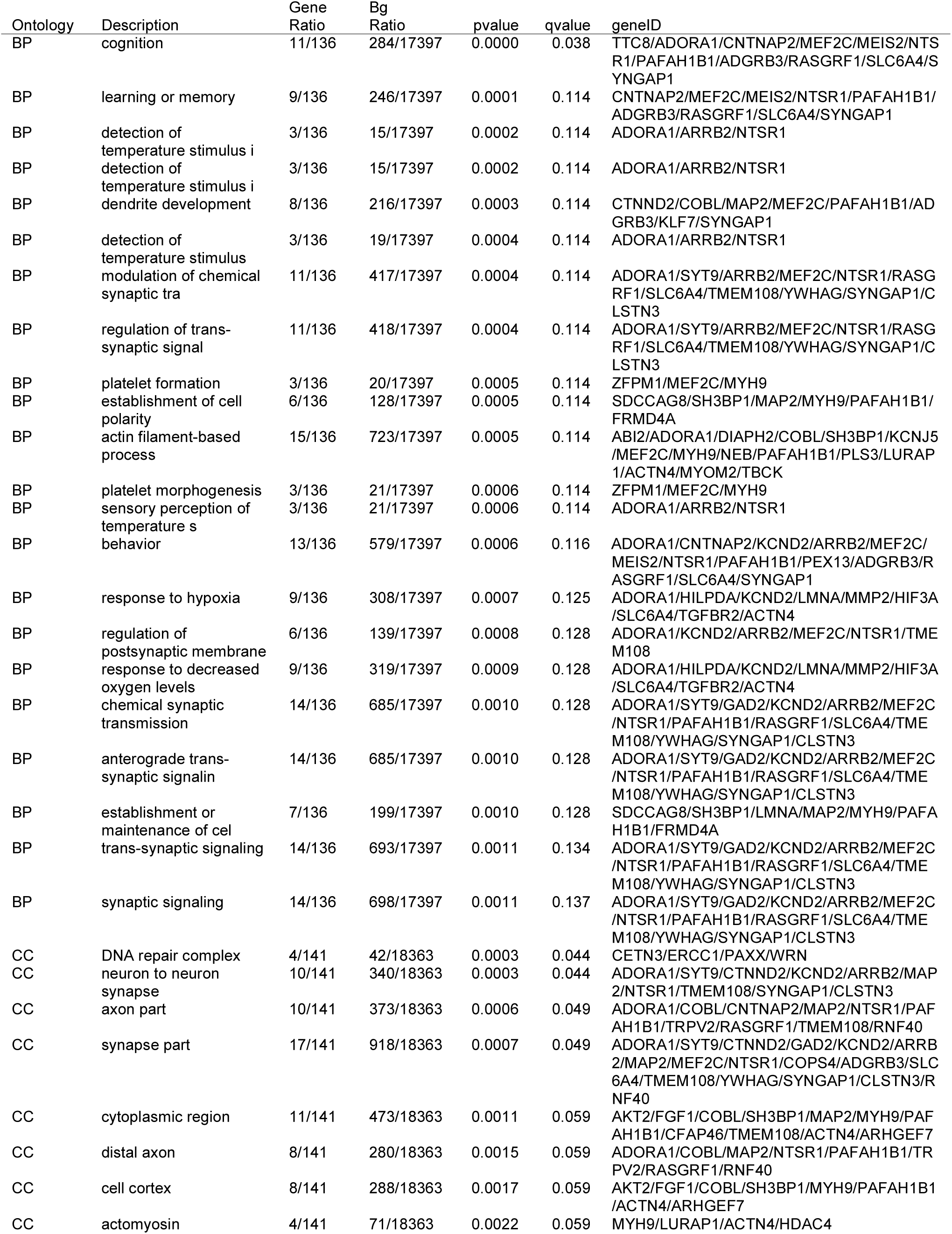
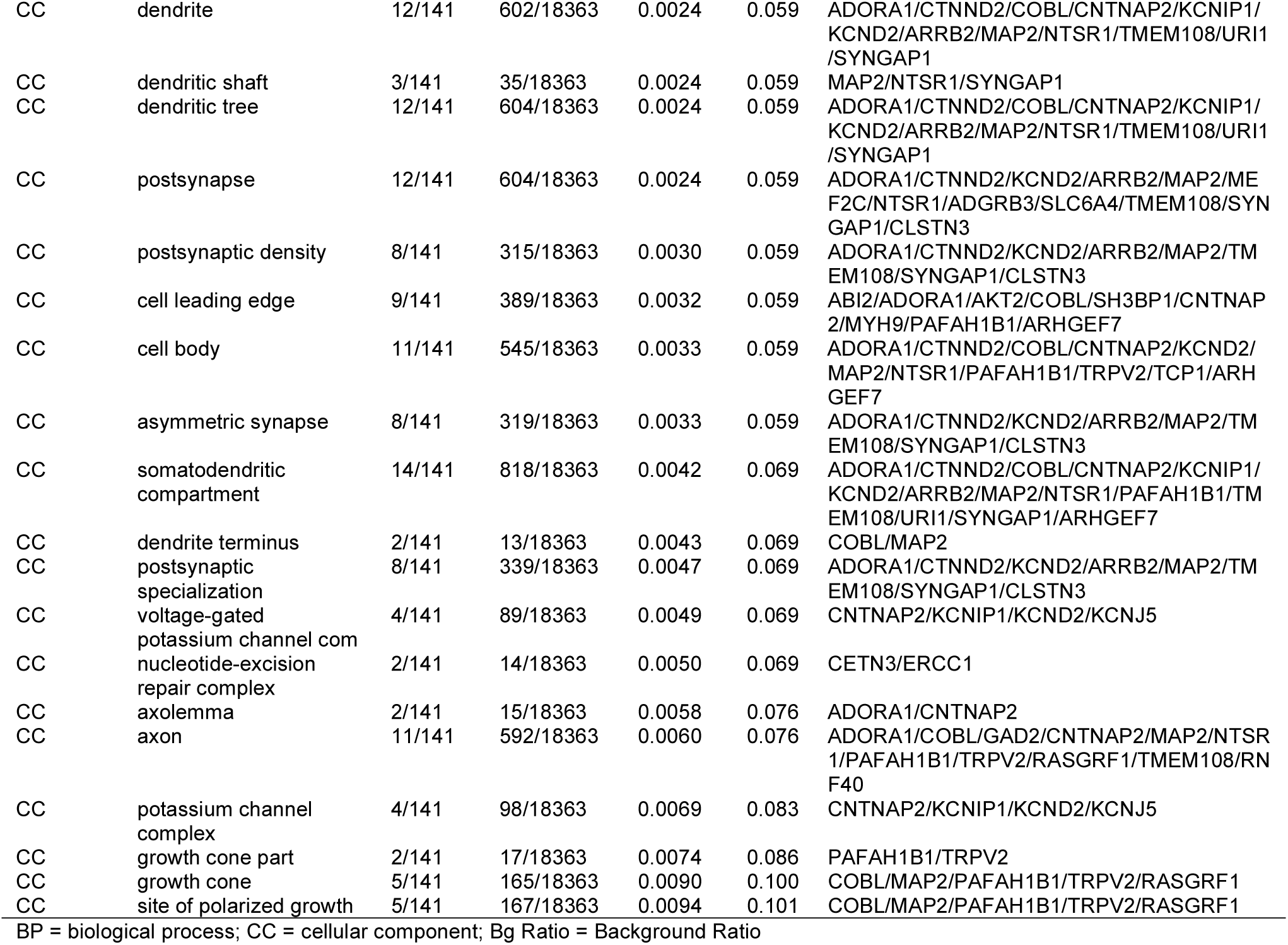
Gene Ontology Results for Differentially Methylated Probes and Regions

Three DMRs overlapped PGC GWAS findings on chromosomes 5, 6, and 16 (p = 0.034; see Table S1 in the supplement for more detail). The overlapping region on chromosome 6 occurred in the major histocompatibility complex (MHC) region and neighbored genomic areas previously associated with early-onset major depression (DNAm)^26^ and depression as defined in the PGC genome-wide allelic meta-analysis (see Figure 1).^54^ None of the sites or regions identified in the AYATS EWAS of early-onset major depression directly overlapped the DMRs associated with postpartum depressive symptom load in this analysis.^26^

**Figure 1.**
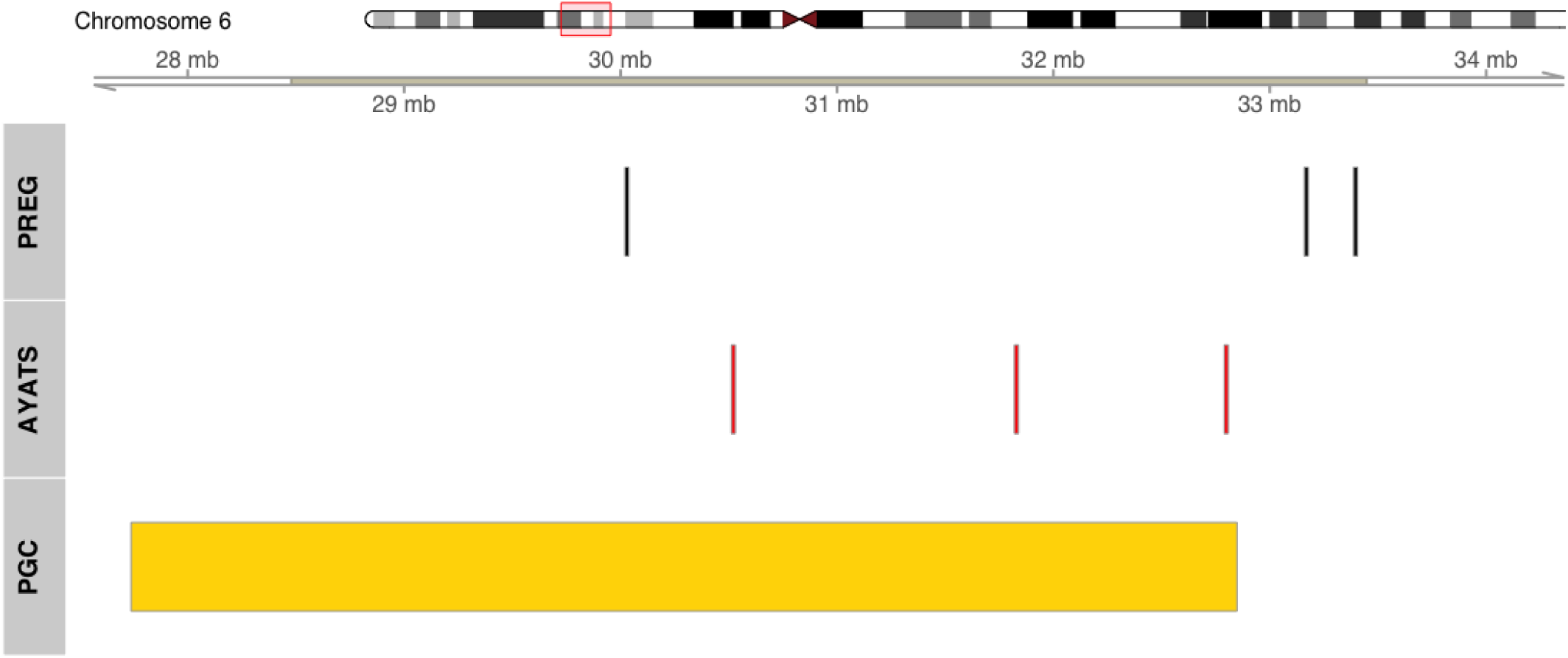
DNA methylation patterns associated with postpartum depressive symptoms (top row) and early-onset major depression (middle row) colocalize to the major histocompatibility complex (MHC) region on chromosome 6. The overlap of the DNA methylation patterns and the genomic region tagged in the genome-wide association meta-analysis of major depression performed by the Psychiatric Genomics Consortium (PGC) is shown in the bottom row (Wray et al. 2018).

**Figure 2.**
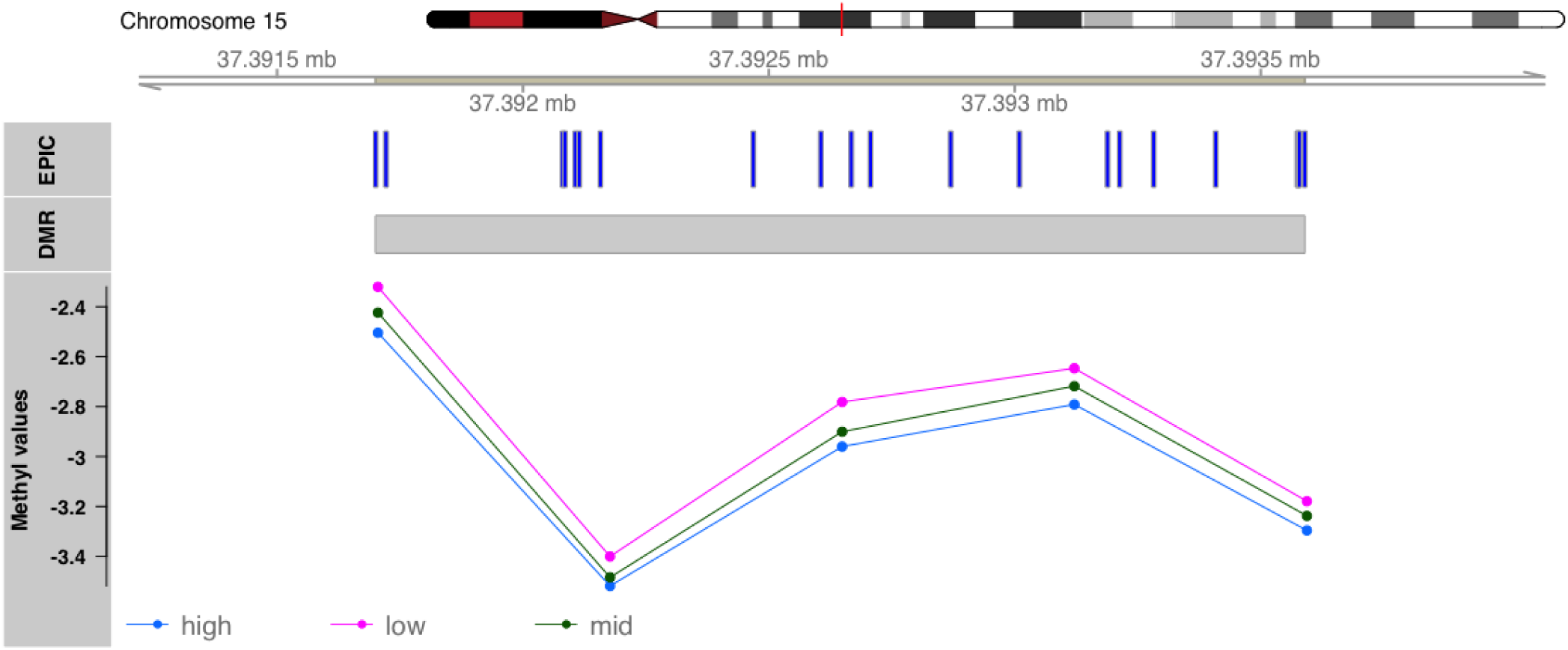
Increases in depressive symptom load negatively correlate with DNA methylation in a significant identified on chromosome 15. The significantly differentially methylated region (row 2, “DMR”) was built using 5 of the 21 CpG probes available on the EPIC array (row 1, “EPIC”). The mean group methyl values are shown for low, mid, and high depressive symptom loads based on Edinburgh Postnatal Depression Scale (EPDS) total score (low = 0-4; mid = 5-12; high = 13 and greater). The threshold of 13 or greater for the high group was selected based on the validated cut-off score for English-speaking women in the postpartum.^31^

## Discussion

Evaluating the credibility and replicability of the significant findings from this analysis is paramount but complicated. Few directly comparable studies exist, and standardized practices for conducting and reporting results from epigenome-wide association studies (EWAS) have not been established. That said, domains to assess the credibility of EWAS results have been proposed, including the level of statistical significance, genomic location, biological relevance, functional relevance, validation of significant associations, and the potential for study design bias or confounding variables to influence the analysis.^29^ An indirect effect of evaluating the results with these criteria is that it also highlights a study’s strengths and weaknesses.

This study identified 98 DMRs and 50 DMPs using DNAm measures from the EPIC beadchip (850k), one of the most comprehensive microarray technologies available to assay DNAm. The combination strategy of site- and region-based analysis identified significant results that overlap a gene set enriched for biological pathways highly relevant to putative depression etiology (e.g., synaptic signaling, dendrite development). Moreover, the significant genome regions include a number of genes that have been previously associated with MD (e.g., *SLC6A4*, also known as *5-HTT* which encodes SERT^56^ and *RNF145*^57^) or with estrogen and progesterone signaling (e.g., *FOXA1, ARRB2, ITGB3BP*), which may be particularly relevant for MDP and perinatal depressive symptom risk liability.^37^ None of the sites or regions from this directly overlapped regions significant in the AYATS EWAS of early-onset MD;^26^ however, both this study and the AYATS EWAS identified significant DNAm regions in the major histocompatibility complex (MHC) region on chromosome 6, which was also identified in the latest PGC GWAS of depression. The two DNAm studies also shared a large proportion of Gene Ontology (GO) terms in their respective gene set enrichment analyses (e.g., synaptic transmission, central nervous system development), suggesting that even if the exact sites differed by study, the biological pathways with associated genes did not.

Comparing these results within the DNAm-MDP literature is difficult because a majority of the studies used candidate gene approaches, which perform best as methods to refine results from genome-wide analyses. The only other genome-wide DNAm-MDP study that used maternal tissue measured prenatal DNAm with the HumanMethylation450 beadchip (450k). No significant results were found, possibly due to a modest sample size (n=38 antenatal maternal blood samples) ^21^ and using only single probe approach. Modest sample sizes are common in EWAS because of technology costs, but regional analyses can mitigate power issues from small sample sizes by reducing the multiple test burden. Ong and Holbrook estimated that using their regional approach, a two group 450k study with 38 people (n=19 per group) would have 61% power to identify results with an effect size of 2 using a regional approach compared to 39% with a single probe analysis.^30^ Another benefit of regional analyses is that the results are more likely to replicate^30^ in part because significant regional results require multiple nearby probes each to have test statistics greater than a chosen threshold and to exhibit the same direction of effect. Together, the use of genome-wide DNAm, a permutation rank-based approach for assessing significance, and probes with test statistics either in the upper or lower 2.5 percentile to test for DMRs each positively influence the credibility of the results. Further, the genomic locations of the significant DMRs increases the credibility of the results as they overlap genes and genomic regions that either directly corroborate previous findings in the literature or that participate in biological pathways hypothesized to be important for depression risk or onset.

Another strength of this study concerns its design. Significant care was taken to minimize potential biases and confounders from biological, behavioral, and technical factors. The women in this study completed multiple comprehensive questionnaires about perinatal health and behaviors, accessed prenatal care relatively early in gestation, and were generally healthy (e.g., no diabetes),^28^ had healthy singleton pregnancies, and completed a postpartum study visit within seven months of delivery. These study design aspects allow variation from behavioral, biological, and technical factors to be measured and accounted for (e.g., cell type heterogeneity, tobacco use, slide and positional effects, signal quality). Additionally, ancestrally-relevant principal components were incorporated to reduce the likelihood of detecting artifacts due to potential population stratification. Visualization of methyl values by self-reported race in significant DMRs suggested that DNAm values did not differ markedly between groups (Figure S1).

Branching out to other depression phenotypes provides both more literature for comparison, but also more uncertainty. For example, it is not immediately clear if the DNAm patterns associated with postpartum depressive symptoms should resemble those associated with other depression phenotypes, including clinical major depression. Though both are depression phenotypes, depressive symptoms and clinical depression are not equivalent.^58–61^ The issue of nonequivalence emerged when comparing the findings in this analysis to published results. For example, Numata et al. reported a significant relationship between the DNAm at cg14472315 and MD case status.^25^ No significant relationship existed between that probe and self-reported postpartum depressive symptoms in this study; however, a nominally significant relationship between that probe and lifetime history of MD was present in this study (tested *post hoc*), reaffirming the difficulty of comparing studies with related but non-identical depressive phenotypes. Another issue with cross-study comparisons of depressive phenotypes is that the genetic factors contributing to risk liability may not be shared completely between sexes.^62,63^ This limitation is trivial for maternal perinatal mental health phenotypes, but crucial when comparing to other studies that included male and female participants.^64^

In a similar vein, the timing of MD episodes or symptoms may reflect different etiologies. Allelic studies of MD suggest that genetic factors influencing risk vary by age of first MD episode onset such that individuals who have earlier onsets have higher polygenic risk scores for schizophrenia and bipolar disorder compared to individuals who have their first MD episodes later in life.^65^ Furthermore, different etiologies may be present within MDP depending on whether symptoms onset prenatally or in the postpartum.^66^ At least one group has investigated the relationship between risk for postpartum depression and perturbed prenatal DNAm patterns in proximity to genomic regions differentially methylated by estrogen.^67^

For all of its strengths, this study was not without limitations. First, the sample size was modest, which limits the statistical power to conduct robust single site analysis. The sample also included relatively few women with severe levels of postpartum depressive symptoms, which may have unique DNAm signatures compared to clinical MDP. Moreover, this study could not determine if the DNAm patterns identified in this study were a consequence of previous episodes of MD (which is a risk factor for perinatal depression^46^) or if they are related to other genetic or environmental factors. Second, detailed information about medication history was not available. Third, the newness of the EPIC microarray means that the probe set has been less well-vetted for cross-hybridization compared to its predecessor.^42,68^ Four, regional analyses are inherently limited because not all probes have the potential to form regions.^30^ This analysis attempted to address that weakness by also performing a single site analysis. Furthermore, the regional analysis algorithm used in this study selected an equal number of probes with positive and negative t values, corresponding to hypermethylation and hypomethylation in cases versus controls. This strategy assumes that equal representation of positive and negative t values will yield the most fair results; however, it is possible that this assumption limited the number of DMRs identified, especially if cases had much more hyper- or hypomethylation. Finally, this study was limited in its assessment of functional relevance. No gene expression, chromatin conformation, or transcription factor binding assays were run concurrently with postpartum DNAm analyses. No algorithms currently exist that can determine the precise change in DNAm necessary to translate into biologically meaningful differences in chromatin shape or gene regulation. As a result, fully understanding the etiology of MDP and depressive symptoms will likely involve integrating repeated measures from multiple biological layers (e.g., genetic sequence, epigenetic mechanisms, transcription, protein),^69,70^ but no single study could measure every biological layer that might be informative about depression etiology, especially not longitudinally.

Another important consideration for interpreting DNAm results is tissue source.^29,71^ It remains unclear how detrimental the use of peripheral blood DNAm is for identifying genomic regions associated with a psychiatric phenotype like depression. On one hand, specific brain regions hold intuitive appeal for MD-DNAm studies, and the cross-tissue similarity between brain and blood appears to be modest and tied to allelic variation;^72^ however, given the well-established link between MD pathophysiology and aberrant immune system functioning,^34–36,73^ peripheral blood may be the best and most feasible option for large or longitudinal studies of stress-related psychiatric traits. Many of the biomarkers associated with MD are transported in the blood (e.g., IL-1, IL-2, IL-6, TNFa, haptoglobin), and some of these immune-related differences appear to persist after depressive episode remission. Not only does that observation fit with epidemiological studies that find individuals with a positive lifetime history of MD remain sensitive to stress and at higher risk for adverse auto-immune and cardiovascular outcomes,^34^ but also it suggests that DNAm patterns detectable in the blood may retain MD-associated differences even after depressive episode remission. Last but not least, peripheral blood may be useful for indexing changes in the relationship between the CNS and immune system. As a sentinel tissue, peripheral blood travels throughout the entire body and can deliver immune cells through the blood-brain barrier. Key players in the CNS like the neurotransmitter serotonin also serve roles in immune-related biological pathways (e.g., leukocyte activation and proliferation, cytokine secretion, chemotaxis, and apoptosis).^36,74^ Serotonin has been linked to a number of psychiatric conditions including MD, and this study identified a significant DMR overlapping the *SLC6A4* gene body, further implicating this genomic region with psychopathology. The ability for the CNS to modulate and respond to signals from the immune system underscores the intimate relationship between brain function and immune system regulation.

The final category Michels et al. (2013) listed as an important factor in establishing the credibility of DNAm associations with phenotypes is validation.^29^ Validating results typically implies either replicating the finding in an independent study (human or animal) or confirming the presence of differentially methylated probes and regions using another technology (e.g., pyrosequencing). While no pyrosequencing was completed, the literature was searched extensively for associations between depressive phenotypes and biological signatures (e.g., allelic, epigenetic, transcription). As previously mentioned, the genomic regions implicated in this study overlap results from the 2018 Psychiatric Genomics Consortium (PGC) genome-wide allelic association study (GWAS) of MD^54^ and are proximal to significant regions and probes from an EWAS of early-onset MD. Network analysis of the 44 significant loci in the PGC meta-analysis implicated biological pathways associated with neural differentiation, synaptic regulation, risk for schizophrenia, immune response, ion-gated channels, and retinoid X receptors.^54^ Similarly, this analysis identified significant DMRs overlapping genes coding for or related to retinoid X receptors (*RXRB, ITGB3BP*), genes integral to adult neurogenesis and synaptic development and positioning (e.g., *AKT2, SYNGAP1, FOXG1, CTNND2, MEF2C, AIMP1*, etc.), risk for schizophrenia (e.g., *CLSTN3, FARSB, MYOM2, SOX2-OT, SLC39A7, SDCCAG8, LRRC36*), immune response (*LRR1, FAM19A2, CMKLR1, MMP2*), ion-gated channels and binding (e.g., *SLC6A4, SLC6A12, SLC39A7, KCNJ5, CLSTN3, PLS3*). While these similarities do not serve as direct replication, they do increase the credibility of the DNAm findings and underscore the potential for DNAm studies to complement GWA studies.

## Conclusions

Future work should take note of the apparent differences in depressive symptoms and clinical MD and seek to refine association studies by limiting phenotypic heterogeneity. For MDP, that means not only accounting for whether the depressive symptoms or episodes onset prenatally versus postnatally, but also disentangling which DNAm patterns are associated with perinatal depressive phenotypes and which reflect pre-pregnancy events of depressive psychopathology. It is possible that women who experience their first instance of MD in the peripartum have a unique DNAm profile compared to those who have recurrent MD and happen to onset during the peripartum. Furthermore, it is unknown whether an episode of MD relatively early in life evokes a persistent perturbation in DNAm patterning that then goes on to affect the risk for additional depressive symptoms and episodes as well as other adverse health outcomes frequently comorbid with depression (e.g., cardiovascular disease and diabetes mellitus). Clarifying the phenotypes associated with DNAm patterns will enable wet lab researchers to characterize the functional and biological relevance of implicated genomic sites and regions, which is an essential step not only for understanding the biological mechanisms associated with risk and resilience to depressive psychopathology but also for developing screening tests and identifying novel pharmacotherapeutic targets.

## Acknowledgements

We are extremely grateful to the families, clinicians, and research staff who participated in the Pregnancy, Race, Environment, Genes (PREG) study and its postpartum extension. Additionally, we would like to thank the Virginia Commonwealth University Epigenetics Journal Club members who provided feedback on the manuscript.

## Ethics approval and consent to participate

Both the PREG study and its postpartum extension were approved by the Virginia Commonwealth Institutional Review Board (#14000), and participants provided written consent for both.

## Authors’ contributions

DML performed the analyses and wrote the first draft of the manuscript. RMK and BTW provided statistical expertise. TPY, PK, and RRN conceived of the analysis, secured the funding, and provided extensive feedback on the manuscript. In addition, TPY wrote the regional analysis script, and RRN provided expertise in psychiatric epidemiology and phenotyping. All authors read the manuscript, gave feedback, and approved the final version.

## Availability of data and materials

The datasets analyzed during the current study are not publicly available due to the nature of the studies’ consent forms and IRB agreements, which allowed participants to opt out of their data being shared due to individual privacy concerns. As a result, restrictions apply to the availability of these data. Individual level data from participants who consented to allow data sharing is available from the corresponding author upon reasonable request.

All summary-level data and methods can be found on the Open Science Framework (https://osf.io/qsc6n).

## List of abbreviations

AYATS: Adolescent and Young Adult Twin Study
CpG: Cytosine-phosphate-guanine
DMP: Differentially methylated probe
DMR: Differentially methylated region
DNAm: DNA methylation
EPDS: Edinburgh Postnatal Depression Scale
EWAS: Epigenome-wide association study
FDR: False discovery rate
GO: Gene ontology
GWAS: Genome-wide association study
MD: Major depression
MDP: Major depression with onset in the peripartum
MHC: Major histocompatibility complex
PCA/ PC: Principal components analysis / Principal component
PGC: Psychiatric Genomics Consortium
PREG: Pregnancy, Race, Environment, Genes study

