## Supplemental File for "DNA Methylation Associated with Postpartum Depressive Symptoms Overlaps Findings from a Genome-wide Association Meta-Analysis of Depression"

**Enrichment testing for overlap with Psychiatric Genomics Consortium (PGC) genome-wide association study meta-analysis of depression**

In order to calculate the 95% confidence interval for the number of observed overlaps between the differentially methylated regions in this study and the PGC, bootstrapping (k=1000 permutations) was used. The frequency that overlaps of 0-10 PGC loci occurred during the permutation analysis can be found in Table S1.

Table S1: Results of bootstrapping confidence intervals for overlap with the PGC GWAS meta-analysis of depression

| Overlaps | 0 | 1 | 2 | 3 | 4 | 5 | 6 | 7 | 8 | 9 | 10 |
|---|---|---|---|---|---|---|---|---|---|---|---|
| Frequency | 57 | 138 | 225 | 228 | 165 | 90 | 58 | 29 | 7 | 1 | 2 |

Frequency = the number of times each overlap occurred in 1000 permutations.
PGC = Psychiatric Genomics Consortium; GWAS = genome-wide association study

**Figure S1.**

Figure S1. Supplement to Figure 2. In the top figure, the ComBat-adjusted methyl values for each participant are shown for each of the five probes used to build the differentially methylated region on chromosome 15. The color indicates self-reported Census-based race category (blue = African-American; red = European-American). The methyl values have been jittered left/right to reduce over plotting, but the height (i.e., methyl value) has not been altered. In the bottom figure, mean-level methyl values for AA and EA participants are shown for each probe.

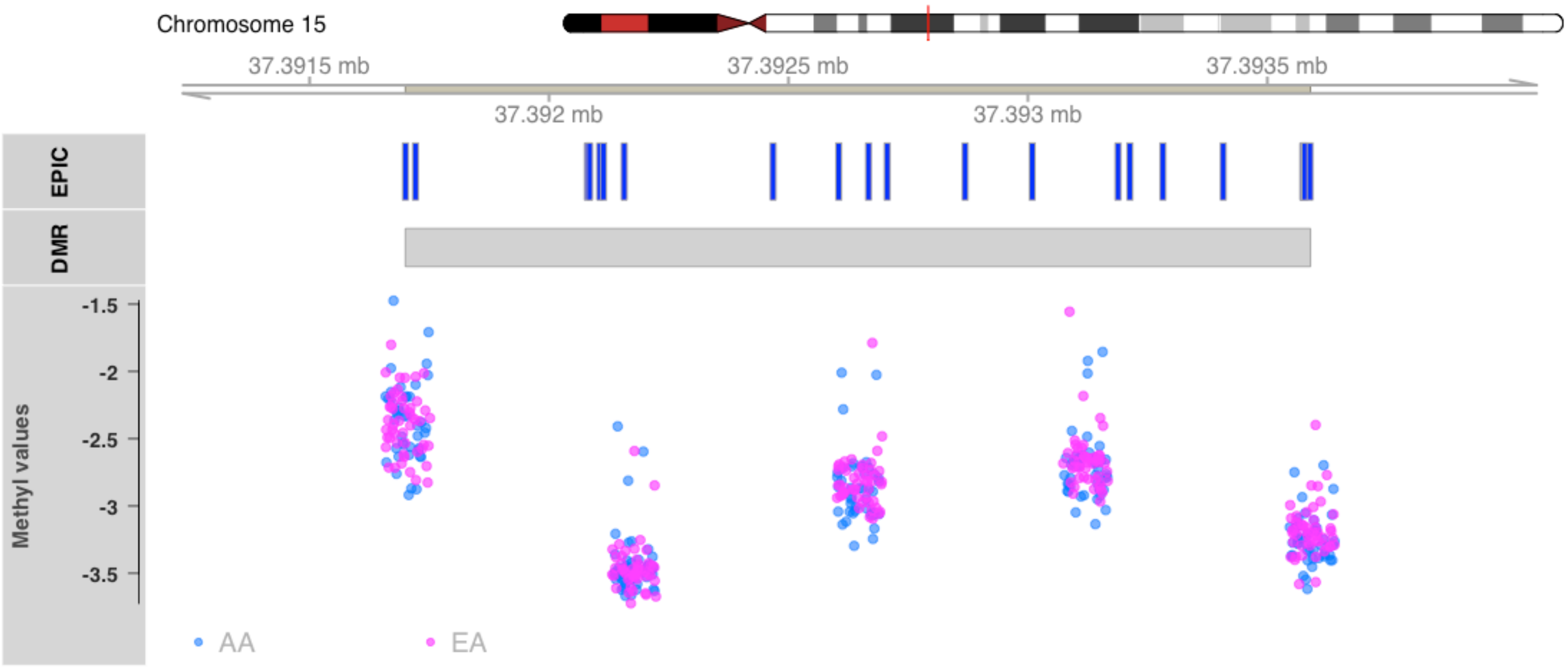

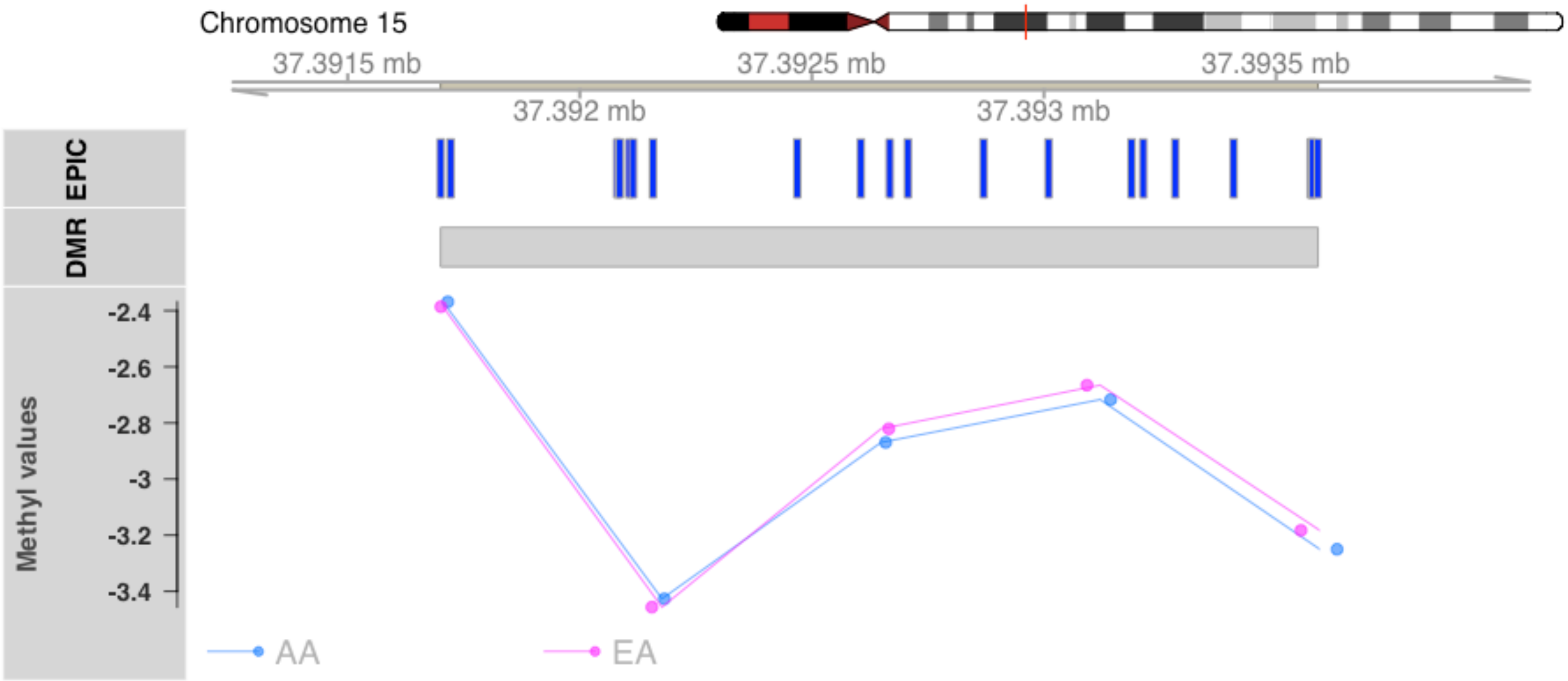

Chromosome 15
37.3915 mb
37.3925 mb
37.3935 mb
37.392 mb
37.393 mb
EPIC
DMR
Methyl values
-2.4
-2.6
-2.8
-3
-3.2
-3.4
AA
EA

### Gene Ontology Supplemental Tables

**DMR & DMP**

| Ontology | Description | GeneRatio | BgRatio | pvalue | qvalue | geneID |
|---|---|---|---|---|---|---|
| BP | cognition | 11/136 | 284/17397 | 0.0000 | 0.038 | TTC8/ADORA1/CNTNAP2/MEF2C/MEIS2/NTSR1/PAFAH1B1/ |
| BP | learning or memory | 9/136 | 246/17397 | 0.0001 | 0.114 | CNTNAP2/MEF2C/MEIS2/NTSR1/PAFAH1B1/ADGRB3/RASGI |
| BP | detection of temperature stimulus i | 3/136 | 15/17397 | 0.0002 | 0.114 | ADORA1/ARRB2/NTSR1 |
| BP | detection of temperature stimulus i | 3/136 | 15/17397 | 0.0002 | 0.114 | ADORA1/ARRB2/NTSR1 |
| BP | dendrite development | 8/136 | 216/17397 | 0.0003 | 0.114 | CTNND2/COBL/MAP2/MEF2C/PAFAH1B1/ADGRB3/KLF7/SY |
| BP | detection of temperature stimulus | 3/136 | 19/17397 | 0.0004 | 0.114 | ADORA1/ARRB2/NTSR1 |
| BP | modulation of chemical synaptic tra | 11/136 | 417/17397 | 0.0004 | 0.114 | ADORA1/SYT9/ARRB2/MEF2C/NTSR1/RASGRF1/SLC6A4/TM |
| BP | regulation of trans-synaptic signal | 11/136 | 418/17397 | 0.0004 | 0.114 | ADORA1/SYT9/ARRB2/MEF2C/NTSR1/RASGRF1/SLC6A4/TM |
| BP | platelet formation | 3/136 | 20/17397 | 0.0005 | 0.114 | ZFPM1/MEF2C/MYH9 |
| BP | establishment of cell polarity | 6/136 | 128/17397 | 0.0005 | 0.114 | SDCCAG8/SH3BP1/MAP2/MYH9/PAFAH1B1/FRMD4A |
| BP | actin filament-based process | 15/136 | 723/17397 | 0.0005 | 0.114 | ABI2/ADORA1/DIAPH2/COBL/SH3BP1/KCNJ5/MEF2C/MYH9 |
| BP | platelet morphogenesis | 3/136 | 21/17397 | 0.0006 | 0.114 | ZFPM1/MEF2C/MYH9 |
| BP | sensory perception of temperature s | 3/136 | 21/17397 | 0.0006 | 0.114 | ADORA1/ARRB2/NTSR1 |
| BP | behavior | 13/136 | 579/17397 | 0.0006 | 0.116 | ADORA1/CNTNAP2/KCND2/ARRB2/MEF2C/MEIS2/NTSR1/PA |
| BP | response to hypoxia | 9/136 | 308/17397 | 0.0007 | 0.125 | ADORA1/HILPDA/KCND2/LMNA/MMP2/HIF3A/SLC6A4/TGF |
| BP | regulation of postsynaptic membrane | 6/136 | 139/17397 | 0.0008 | 0.128 | ADORA1/KCND2/ARRB2/MEF2C/NTSR1/TMEM108 |
| BP | response to decreased oxygen levels | 9/136 | 319/17397 | 0.0009 | 0.128 | ADORA1/HILPDA/KCND2/LMNA/MMP2/HIF3A/SLC6A4/TGF |
| BP | chemical synaptic transmission | 14/136 | 685/17397 | 0.0010 | 0.128 | ADORA1/SYT9/GAD2/KCND2/ARRB2/MEF2C/NTSR1/PAFAH |
| BP | anterograde trans-synaptic signalin | 14/136 | 685/17397 | 0.0010 | 0.128 | ADORA1/SYT9/GAD2/KCND2/ARRB2/MEF2C/NTSR1/PAFAH |
| BP | establishment or maintenance of cel | 7/136 | 199/17397 | 0.0010 | 0.128 | SDCCAG8/SH3BP1/LMNA/MAP2/MYH9/PAFAH1B1/FRMD4A |
| BP | trans-synaptic signaling | 14/136 | 693/17397 | 0.0011 | 0.134 | ADORA1/SYT9/GAD2/KCND2/ARRB2/MEF2C/NTSR1/PAFAH |
| BP | synaptic signaling | 14/136 | 698/17397 | 0.0011 | 0.137 | ADORA1/SYT9/GAD2/KCND2/ARRB2/MEF2C/NTSR1/PAFAH |
| BP | striated muscle cell differentiatio | 8/136 | 269/17397 | 0.0013 | 0.140 | LMNA/ARRB2/MEF2C/MYH9/NEB/ADGRB3/MYOM2/HDAC4 |
| BP | actin cytoskeleton organization | 13/136 | 633/17397 | 0.0014 | 0.140 | ABI2/DIAPH2/COBL/SH3BP1/MEF2C/MYH9/NEB/PAFAH1B1/ |
| BP | muscle hypertrophy in response to s | 3/136 | 29/17397 | 0.0015 | 0.140 | LMNA/MEF2C/HDAC4 |
| BP | cardiac muscle adaptation | 3/136 | 29/17397 | 0.0015 | 0.140 | LMNA/MEF2C/HDAC4 |
| BP | cardiac muscle hypertrophy in respo | 3/136 | 29/17397 | 0.0015 | 0.140 | LMNA/MEF2C/HDAC4 |
| BP | response to oxygen levels | 9/136 | 342/17397 | 0.0015 | 0.140 | ADORA1/HILPDA/KCND2/LMNA/MMP2/HIF3A/SLC6A4/TGF |
| BP | cardiac muscle tissue development | 7/136 | 216/17397 | 0.0016 | 0.142 | ZFPM1/LMNA/ARRB2/MEF2C/NEB/TGFBR2/MYOM2 |
| BP | supramolecular fiber organization | 13/136 | 645/17397 | 0.0016 | 0.144 | ABI2/CETN3/DIAPH2/COBL/SH3BP1/FIGNL2/MAP2/MEF2C/ |
| CC | DNA repair complex | 4/141 | 42/18363 | 0.0003 | 0.044 | CETN3/ERCC1/PAXX/WRN |
| CC | neuron to neuron synapse | 10/141 | 340/18363 | 0.0003 | 0.044 | ADORA1/SYT9/CTNND2/KCND2/ARRB2/MAP2/NTSR1/TMEI |

| Ontology | Description | GeneRatio | BgRatio | pvalue | qvalue | geneID |
|---|---|---|---|---|---|---|
| CC | axon part | 10/141 | 373/18363 | 0.0006 | 0.049 | ADORA1/COBL/CNTNAP2/MAP2/NTSR1/PAFAH1B1/TRPV2/ |
| CC | synapse part | 17/141 | 918/18363 | 0.0007 | 0.049 | ADORA1/SYT9/CTNND2/GAD2/KCND2/ARRB2/MAP2/MEF2C |
| CC | cytoplasmic region | 11/141 | 473/18363 | 0.0011 | 0.059 | AKT2/FGF1/COBL/SH3BP1/MAP2/MYH9/PAFAH1B1/CFAP46/ |
| CC | distal axon | 8/141 | 280/18363 | 0.0015 | 0.059 | ADORA1/COBL/MAP2/NTSR1/PAFAH1B1/TRPV2/RASGRF1/ |
| CC | cell cortex | 8/141 | 288/18363 | 0.0017 | 0.059 | AKT2/FGF1/COBL/SH3BP1/MYH9/PAFAH1B1/ACTN4/ARHGI |
| CC | actomyosin | 4/141 | 71/18363 | 0.0022 | 0.059 | MYH9/LURAP1/ACTN4/HDAC4 |
| CC | dendrite | 12/141 | 602/18363 | 0.0024 | 0.059 | ADORA1/CTNND2/COBL/CNTNAP2/KCNIP1/KCND2/ARRB2/ |
| CC | dendritic shaft | 3/141 | 35/18363 | 0.0024 | 0.059 | MAP2/NTSR1/SYNGAP1 |
| CC | dendritic tree | 12/141 | 604/18363 | 0.0024 | 0.059 | ADORA1/CTNND2/COBL/CNTNAP2/KCNIP1/KCND2/ARRB2/ |
| CC | postsynapse | 12/141 | 604/18363 | 0.0024 | 0.059 | ADORA1/CTNND2/KCND2/ARRB2/MAP2/MEF2C/NTSR1/AD |
| CC | postsynaptic density | 8/141 | 315/18363 | 0.0030 | 0.059 | ADORA1/CTNND2/KCND2/ARRB2/MAP2/TMEM108/SYNGAP |
| CC | cell leading edge | 9/141 | 389/18363 | 0.0032 | 0.059 | ABI2/ADORA1/AKT2/COBL/SH3BP1/CNTNAP2/MYH9/PAFAI |
| CC | cell body | 11/141 | 545/18363 | 0.0033 | 0.059 | ADORA1/CTNND2/COBL/CNTNAP2/KCND2/MAP2/NTSR1/PA |
| CC | asymmetric synapse | 8/141 | 319/18363 | 0.0033 | 0.059 | ADORA1/CTNND2/KCND2/ARRB2/MAP2/TMEM108/SYNGAP |
| CC | somatodendritic compartment | 14/141 | 818/18363 | 0.0042 | 0.069 | ADORA1/CTNND2/COBL/CNTNAP2/KCNIP1/KCND2/ARRB2/ |
| CC | dendrite terminus | 2/141 | 13/18363 | 0.0043 | 0.069 | COBL/MAP2 |
| CC | postsynaptic specialization | 8/141 | 339/18363 | 0.0047 | 0.069 | ADORA1/CTNND2/KCND2/ARRB2/MAP2/TMEM108/SYNGAP |
| CC | voltage-gated potassium channel com | 4/141 | 89/18363 | 0.0049 | 0.069 | CNTNAP2/KCNIP1/KCND2/KCNJ5 |
| CC | nucleotide-excision repair complex | 2/141 | 14/18363 | 0.0050 | 0.069 | CETN3/ERCC1 |
| CC | axolemma | 2/141 | 15/18363 | 0.0058 | 0.076 | ADORA1/CNTNAP2 |
| CC | axon | 11/141 | 592/18363 | 0.0060 | 0.076 | ADORA1/COBL/GAD2/CNTNAP2/MAP2/NTSR1/PAFAH1B1/T |
| CC | potassium channel complex | 4/141 | 98/18363 | 0.0069 | 0.083 | CNTNAP2/KCNIP1/KCND2/KCNJ5 |
| CC | growth cone part | 2/141 | 17/18363 | 0.0074 | 0.086 | PAFAH1B1/TRPV2 |
| CC | growth cone | 5/141 | 165/18363 | 0.0090 | 0.100 | COBL/MAP2/PAFAH1B1/TRPV2/RASGRF1 |
| CC | site of polarized growth | 5/141 | 167/18363 | 0.0094 | 0.101 | COBL/MAP2/PAFAH1B1/TRPV2/RASGRF1 |

**DMR only**

| Ontology | Description | GeneRatio | BgRatio | pvalue | qvalue | geneID |
|---|---|---|---|---|---|---|
| BP | platelet formation | 3/97 | 20/17137 | 0.0002 | 0.232 | ZFPM1/MEF2C/MYH9 |
| BP | platelet morphogenesis | 3/97 | 21/17137 | 0.0002 | 0.232 | ZFPM1/MEF2C/MYH9 |
| BP | cardiac muscle tissue development | 6/97 | 214/17137 | 0.0014 | 0.410 | ZFPM1/LMNA/ARRB2/MEF2C/TGFBR2/MYOM2 |
| BP | regulation of macrophage apoptoti | 2/97 | 10/17137 | 0.0014 | 0.410 | MEF2C/TCP1 |
| BP | macrophage apoptotic process | 2/97 | 12/17137 | 0.0020 | 0.410 | MEF2C/TCP1 |
| BP | heart valve morphogenesis | 3/97 | 47/17137 | 0.0024 | 0.410 | ZFPM1/MEF2C/TGFBR2 |
| BP | actin filament-based process | 11/97 | 721/17137 | 0.0025 | 0.410 | ABI2/ADORA1/DIAPH2/SH3BP1/KCNJ5/MEF2C/MYH9/PLS3/L |

| Ontology | Description | GeneRatio | BgRatio | pvalue | qvalue | geneID |
|---|---|---|---|---|---|---|
| BP | excitatory postsynaptic potential | 4/97 | 103/17137 | 0.0028 | 0.410 | ADORA1/ARRB2/MEF2C/TMEM108 |
| BP | lung morphogenesis | 3/97 | 50/17137 | 0.0028 | 0.410 | FOXA1/HHIP/TGFBR2 |
| BP | negative regulation of interleuki | 2/97 | 15/17137 | 0.0032 | 0.410 | CMKLR1/ARRB2 |
| BP | detection of temperature stimulus | 2/97 | 15/17137 | 0.0032 | 0.410 | ADORA1/ARRB2 |
| BP | detection of temperature stimulus | 2/97 | 15/17137 | 0.0032 | 0.410 | ADORA1/ARRB2 |
| BP | heart valve development | 3/97 | 53/17137 | 0.0034 | 0.410 | ZFPM1/MEF2C/TGFBR2 |
| BP | regulation of synaptic transmissi | 2/97 | 16/17137 | 0.0036 | 0.410 | ARRB2/SLC6A4 |
| BP | chemical synaptic transmission, p | 4/97 | 111/17137 | 0.0036 | 0.410 | ADORA1/ARRB2/MEF2C/TMEM108 |
| BP | positive regulation of intracellu | 2/97 | 17/17137 | 0.0041 | 0.410 | FOXA1/LMO3 |
| BP | ventricular cardiac muscle cell d | 2/97 | 17/17137 | 0.0041 | 0.410 | LMNA/MEF2C |
| BP | striated muscle cell differentiat | 6/97 | 268/17137 | 0.0042 | 0.410 | LMNA/ARRB2/MEF2C/MYH9/ADGRB3/MYOM2 |
| BP | striated muscle tissue developmen | 7/97 | 367/17137 | 0.0048 | 0.410 | ZFPM1/LMNA/ARRB2/MEF2C/HIVEP3/TGFBR2/MYOM2 |
| BP | animal organ formation | 3/97 | 61/17137 | 0.0050 | 0.410 | FGF1/MEF2C/TGFBR2 |
| BP | detection of temperature stimulus | 2/97 | 19/17137 | 0.0051 | 0.410 | ADORA1/ARRB2 |
| BP | negative regulation of release of | 2/97 | 19/17137 | 0.0051 | 0.410 | LMNA/ARRB2 |
| BP | cardiac muscle cell differentiati | 4/97 | 123/17137 | 0.0052 | 0.410 | LMNA/ARRB2/MEF2C/MYOM2 |
| BP | establishment or maintenance of c | 5/97 | 199/17137 | 0.0055 | 0.410 | SDCCAG8/SH3BP1/LMNA/MYH9/FRMD4A |
| BP | cognition | 6/97 | 284/17137 | 0.0055 | 0.410 | TTC8/ADORA1/MEF2C/MEIS2/ADGRB3/SLC6A4 |
| BP | negative regulation of organic ac | 2/97 | 20/17137 | 0.0056 | 0.410 | ADORA1/AKT2 |
| BP | muscle tissue development | 7/97 | 382/17137 | 0.0060 | 0.410 | ZFPM1/LMNA/ARRB2/MEF2C/HIVEP3/TGFBR2/MYOM2 |
| BP | establishment of cell polarity | 4/97 | 128/17137 | 0.0060 | 0.410 | SDCCAG8/SH3BP1/MYH9/FRMD4A |
| BP | embryonic hemopoiesis | 2/97 | 21/17137 | 0.0062 | 0.410 | ZFPM1/TGFBR2 |
| BP | sensory perception of temperature | 2/97 | 21/17137 | 0.0062 | 0.410 | ADORA1/ARRB2 |
| BP | atrioventricular valve morphogene | 2/97 | 22/17137 | 0.0068 | 0.410 | ZFPM1/TGFBR2 |
| BP | inflammatory cell apoptotic proce | 2/97 | 22/17137 | 0.0068 | 0.410 | MEF2C/TCP1 |
| BP | negative regulation of endothelia | 3/97 | 69/17137 | 0.0070 | 0.410 | MIR129-2/MEF2C/AIMP1 |
| BP | positive regulation of lipid cata | 2/97 | 23/17137 | 0.0074 | 0.410 | ADORA1/AKT2 |
| BP | tissue morphogenesis | 9/97 | 608/17137 | 0.0075 | 0.410 | ZFPM1/FGF1/SH3BP1/FOXA1/ARRB2/MEF2C/HHIP/TGFBR2/[illegible] |
| BP | regulation of postsynaptic membra | 4/97 | 139/17137 | 0.0080 | 0.410 | ADORA1/ARRB2/MEF2C/TMEM108 |
| BP | atrioventricular valve developmen | 2/97 | 24/17137 | 0.0081 | 0.410 | ZFPM1/TGFBR2 |
| BP | negative regulation of vascular e | 2/97 | 24/17137 | 0.0081 | 0.410 | MIR129-2/MEF2C |
| BP | outflow tract morphogenesis | 3/97 | 76/17137 | 0.0092 | 0.410 | ZFPM1/MEF2C/TGFBR2 |
| BP | negative regulation of anion tran | 2/97 | 26/17137 | 0.0094 | 0.410 | ADORA1/AKT2 |
| BP | actin cytoskeleton organization | 9/97 | 631/17137 | 0.0095 | 0.410 | ABI2/DIAPH2/SH3BP1/MEF2C/MYH9/PLS3/LURAP1/MYOM2/[illegible] |
| MF | ATP-dependent helicase activity | 3/93 | 73/16598 | 0.0080 | 0.402 | CHD3/WRN/DHX38 |
| MF | purine NTP-dependent helicase act | 3/93 | 73/16598 | 0.0080 | 0.402 | CHD3/WRN/DHX38 |
| MF | neurotransmitter:sodium symporter | 2/93 | 27/16598 | 0.0100 | 0.402 | SLC6A4/SLC6A12 |

| Ontology | Description | GeneRatio | BgRatio | pvalue | qvalue | geneID |
|---|---|---|---|---|---|---|
| CC | DNA repair complex | 3/100 | 41/18080 | 0.0015 | 0.443 | CETN3/PAXX/WRN |
| CC | myosin filament | 2/100 | 22/18080 | 0.0065 | 0.475 | MYH9/MYOM2 |

**DMP only**

| Ontology | Description | GeneRatio | BgRatio | pvalue | qvalue | geneID |
|---|---|---|---|---|---|---|
| BP | central nervous system neuron dev | 3/38 | 69/16678 | 0.0005 | 0.238 | DCLK2/MAP2/PAFAH1B1 |
| BP | transmission of nerve impulse | 3/38 | 71/16678 | 0.0006 | 0.238 | CNTNAP2/KCND2/PAFAH1B1 |
| BP | central nervous system neuron dif | 4/38 | 170/16678 | 0.0006 | 0.238 | DCLK2/MAP2/PAFAH1B1/WNT9B |
| BP | multicellular organismal signalin | 4/38 | 199/16678 | 0.0011 | 0.245 | CNTNAP2/KCNIP1/KCND2/PAFAH1B1 |
| BP | limbic system development | 3/38 | 101/16678 | 0.0016 | 0.245 | DCLK2/CNTNAP2/PAFAH1B1 |
| BP | learning or memory | 4/38 | 233/16678 | 0.0019 | 0.245 | CNTNAP2/NTSR1/PAFAH1B1/RASGRF1 |
| BP | chemical synaptic transmission | 6/38 | 603/16678 | 0.0022 | 0.245 | SYT9/KCND2/NTSR1/PAFAH1B1/RASGRF1/YWHAG |
| BP | anterograde trans-synaptic signal | 6/38 | 603/16678 | 0.0022 | 0.245 | SYT9/KCND2/NTSR1/PAFAH1B1/RASGRF1/YWHAG |
| BP | synaptic signaling | 6/38 | 605/16678 | 0.0023 | 0.245 | SYT9/KCND2/NTSR1/PAFAH1B1/RASGRF1/YWHAG |
| BP | trans-synaptic signaling | 6/38 | 605/16678 | 0.0023 | 0.245 | SYT9/KCND2/NTSR1/PAFAH1B1/RASGRF1/YWHAG |
| BP | establishment of cell polarity | 3/38 | 120/16678 | 0.0026 | 0.245 | MAP2/PAFAH1B1/FRMD4A |
| BP | phosphatidylcholine biosynthetic | 2/38 | 36/16678 | 0.0030 | 0.245 | PEMT/CHKA |
| BP | regulation of ion transmembrane t | 5/38 | 440/16678 | 0.0031 | 0.245 | KCNIP1/KCND2/NTSR1/RASGRF1/ACTN4 |
| BP | cognition | 4/38 | 270/16678 | 0.0032 | 0.245 | CNTNAP2/NTSR1/PAFAH1B1/RASGRF1 |
| BP | regulation of transmembrane trans | 5/38 | 455/16678 | 0.0035 | 0.245 | KCNIP1/KCND2/NTSR1/RASGRF1/ACTN4 |
| BP | adult behavior | 3/38 | 140/16678 | 0.0039 | 0.245 | CNTNAP2/NTSR1/PAFAH1B1 |
| BP | regulation of cation transmembran | 4/38 | 287/16678 | 0.0040 | 0.245 | KCNIP1/NTSR1/RASGRF1/ACTN4 |
| BP | vesicle cytoskeletal trafficking | 2/38 | 43/16678 | 0.0043 | 0.245 | PAFAH1B1/ACTN4 |
| BP | pallium development | 3/38 | 164/16678 | 0.0061 | 0.245 | DCLK2/CNTNAP2/PAFAH1B1 |
| BP | establishment or maintenance of c | 3/38 | 182/16678 | 0.0081 | 0.245 | MAP2/PAFAH1B1/FRMD4A |
| BP | behavior | 5/38 | 574/16678 | 0.0093 | 0.245 | CNTNAP2/KCND2/NTSR1/PAFAH1B1/RASGRF1 |
| BP | locomotory behavior | 3/38 | 192/16678 | 0.0094 | 0.245 | KCND2/NTSR1/PAFAH1B1 |
| BP | ammonium ion metabolic process | 3/38 | 194/16678 | 0.0097 | 0.245 | PEMT/CHKA/PAFAH1B1 |
| BP | positive regulation of dendrite d | 2/38 | 66/16678 | 0.0099 | 0.245 | COBL/PAFAH1B1 |
| CC | dendrite | 7/40 | 492/17666 | 0.0001 | 0.009 | COBL/CNTNAP2/KCNIP1/KCND2/MAP2/NTSR1/URI1 |
| CC | somatodendritic compartment | 8/40 | 690/17666 | 0.0001 | 0.009 | COBL/CNTNAP2/KCNIP1/KCND2/MAP2/NTSR1/PAFAH1B1/UR |
| CC | neuronal cell body | 6/40 | 435/17666 | 0.0004 | 0.014 | COBL/CNTNAP2/KCND2/MAP2/NTSR1/PAFAH1B1 |
| CC | axon part | 4/40 | 180/17666 | 0.0007 | 0.014 | COBL/CNTNAP2/NTSR1/PAFAH1B1 |
| CC | supramolecular fiber | 8/40 | 894/17666 | 0.0008 | 0.014 | DCLK2/COBL/KRTAP19-4/MAP2/NEB/PAFAH1B1/ACTN4/HDAC |
| CC | supramolecular polymer | 8/40 | 901/17666 | 0.0008 | 0.014 | DCLK2/COBL/KRTAP19-4/MAP2/NEB/PAFAH1B1/ACTN4/HDAC |

| Ontology | Description | GeneRatio | BgRatio | pvalue | qvalue | geneID |
|---|---|---|---|---|---|---|
| CC | supramolecular complex | 8/40 | 902/17666 | 0.0008 | 0.014 | DCLK2/COBL/KRTAP19-4/MAP2/NEB/PAFAH1B1/ACTN4/HDAC |
| CC | cell body | 6/40 | 499/17666 | 0.0008 | 0.014 | COBL/CNTNAP2/KCND2/MAP2/NTSR1/PAFAH1B1 |
| CC | voltage-gated potassium channel c | 3/40 | 94/17666 | 0.0012 | 0.019 | CNTNAP2/KCNIP1/KCND2 |
| CC | potassium channel complex | 3/40 | 98/17666 | 0.0014 | 0.019 | CNTNAP2/KCNIP1/KCND2 |
| CC | perikaryon | 3/40 | 114/17666 | 0.0022 | 0.027 | CNTNAP2/KCND2/NTSR1 |
| CC | Z disc | 3/40 | 119/17666 | 0.0025 | 0.028 | NEB/ACTN4/HDAC4 |
| CC | dendritic shaft | 2/40 | 36/17666 | 0.0030 | 0.031 | MAP2/NTSR1 |
| CC | I band | 3/40 | 132/17666 | 0.0033 | 0.032 | NEB/ACTN4/HDAC4 |
| CC | cluster of actin-based cell proje | 3/40 | 140/17666 | 0.0039 | 0.035 | PEMT/PAFAH1B1/ACTN4 |
| CC | growth cone | 3/40 | 151/17666 | 0.0048 | 0.040 | COBL/PAFAH1B1/RASGRF1 |
| CC | site of polarized growth | 3/40 | 156/17666 | 0.0053 | 0.042 | COBL/PAFAH1B1/RASGRF1 |
| CC | actomyosin | 2/40 | 64/17666 | 0.0092 | 0.066 | ACTN4/HDAC4 |
| CC | sarcomere | 3/40 | 192/17666 | 0.0093 | 0.066 | NEB/ACTN4/HDAC4 |
